# Morphogenic, molecular, and cellular adaptations for unidirectional airflow in the chicken lung

**DOI:** 10.1101/2024.08.20.608866

**Authors:** Kamryn N Gerner-Mauro, Lisandra Vila Ellis, Guolun Wang, Richa Nayak, Peter Y Lwigale, Ross A Poché, Jichao Chen

## Abstract

Unidirectional airflow in the avian lung enables gas exchange during both inhalation and exhalation. The underlying developmental process and how it deviates from that of the bidirectional mammalian lung are poorly understood. Sampling key developmental stages with multiscale 3D imaging and single-cell transcriptomics, we delineate morphogenic, molecular, and cellular features that accommodate the unidirectional airflow in the chicken lung. Primary termini of hyper-elongated branches are eliminated via proximal-short and distal-long fusions, forming parabronchi. Neoform termini extend radially through parabronchial smooth muscle to form gas-exchanging alveoli. Supporting this radial alveologenesis, branch stalks halt their proximalization, defined by SOX9-SOX2 transition, and become SOX9^low^ parabronchi. Primary and secondary vascular plexi interface with primary and neoform termini, respectively. Single-cell and Stereo-seq spatial transcriptomics reveal a third, chicken-specific alveolar cell type expressing KRT14, hereby named luminal cells. Luminal, alveolar type 2, and alveolar type 1 cells sequentially occupy concentric zones radiating from the parabronchial lumen. Our study explores the evolutionary space of lung diversification and lays the foundation for functional analysis of species-specific genetic determinants.

## INTRODUCTION

The engineering challenge to boost the gas exchange capacity of the lung is met with an increase in complexity and efficiency. Swim bladders are postulated to be repurposed from buoyancy control to gas exchange because they (1) hold secreted rather than inhaled gases, (2) have anti-collapsing surfactant coating, and (3) share a common foregut origin with the lung, while exhibiting mutual exclusivity in a given animal, especially lungfishes (Maina, 1987; Perry *et al*., 2001; Pelster, 2021; Werneburg, Hoßfeld and Levit, 2024). The surface area of the gas exchange interface expands substantially by forming inner folding and secondary septation in amphibian and mammalian lungs, respectively, and exponentially through branching morphogenesis to aerate millions of alveoli (Vila Ellis and Chen, 2021). Besides increasing structural complexity, the physics of gas diffusion calls for a thinner diffusion barrier, as implemented in the basement membrane-sharing arrangement of the ultrathin epithelium and endothelium, and a steeper oxygen gradient (Hsia, Hyde and Weibel, 2016). The latter seems a constant tied to atmospheric oxygen concentrations but varies between inhalation and exhalation in the bidirectional mammalian lung. By enabling unidirectional airflow with a continuous supply of fresh air and gas exchange during exhalation, the avian lung improves the net oxygen gradient averaged over time (Brackenbury, 1971; Kuethe, 1988; O’Connor and Claessens, 2005).

The more efficient unidirectional design poses structural mandates on lung morphogenesis. First, airflow is still bidirectional in the avian trachea, so part of inhaled and exhaled air needs to be temporarily stored while the rest passes through the lung. Accordingly, multiple air sacs form proximal and distal to the lung and extend into bones to also lighten them for flight (Wedel, 2003; O’Connor, 2004). Second, the remaining primary termini from branching morphogenesis fuse to form continuous passages called parabronchi (Duncker, 1972). Consequently, the cul-de-sac branch tips that will give rise to the alveoli of the mammalian lung are eliminated, coincidentally bypassing the need for extensive branching. Instead, neoform protrusions grow radially from the parabronchi to form the alveoli with air passing sequentially through acini, infundibula, and air capillaries (Maina, 2022).

Such structural constraint demands a fresh look into underlying molecular and cellular events that have been ingrained through studies of the mammalian lung. For example, branch tips and stalks from the initial rounds of branching morphogenesis must disengage from the molecular machinery driving the fractal cycles of budding and bifurcation in the mammalian lung. The two waves of branching and airway maturation, respectively marked by SOX9 and SOX2 (Alanis *et al*., 2014), must also be uncoupled to accommodate the avian-specific parabronchi that differ from the more proximal, airway-equivalent fundamental bronchi. Relatedly, the cell types of the parabronchi and alveoli, including the presence of gas-exchanging alveolar type 1 (AT1) cells, are subject to debate (Klika *et al*., 1996; Watson, Fu and West, 2007, 2007; West *et al*., 2010).

This study seeks to define the morphogenic, molecular, and cellular features throughout embryonic chicken lung development that underlie its unidirectional airflow. Building on classic morphometric and selected molecular analyses, and recent careful documentation of early avian lung development (Maina, 1984, 2002; Sakiyama, Yamagishi and Kuroiwa, 2003; Tzou *et al*., 2016; Spurlin *et al*., 2019; Palmer and Nelson, 2020; Goodwin *et al*., 2022), we use multiscale whole-lung imaging to spatiotemporally map two types of fusion, proximal epithelium specification, and co-development of the vasculature. We also use single-cell transcriptomics and spatial transcriptomics to discover a previously unknown cell type lining the parabronchial lumen, as well as candidate molecules controlling evolutionary innovations.

## RESULTS

### Multiscale time-course 3D imaging identifies proximal-short and distal-long fusions of hyper-elongated primary termini

Despite the remarkable parallel in early lateral budding of the avian lung to the mammalian lung, branch complexity increases exponentially over time, exacerbated by topological changes due to fusion (Metzger *et al*., 2008; Tzou *et al*., 2016; Goodwin and Nelson, 2020). To better document this structural complexity in 3D, we used optical projection tomography (OPT), a mesoscale, isometric imaging method (Sharpe *et al*., 2002), to visualize and quantify epithelial tubulogenesis and the associated smooth muscle in the entire chicken lung up to embryonic day (E) 14 when fusion is complete (Palmer and Nelson, 2020) (Fig. 1A). At E6 and E7, a row of side branches (B1 through B8) emerged along the main bronchus, with rotational offset-like runs in a spiral staircase, unlike the more defined lateral, medial, dorsal, and ventral domains of the mouse lung (Metzger *et al*., 2008) (Fig. 1B). Tips of those side branches dilated and grew multiple new buds (Fig. 1C). Each new bud elongated and widened, seeding the next round of buds (Fig. 1C). This budding cycle was not limited to a single growth plane and thus formed an array, rather than a sheet, of parallel branches (Fig. 1C, 1D asterisks), in contrast to the bifurcation cycle in the mouse lung where branch tip splitting favored a barbed-wire structure (Fig. 1C schematic). Consequently, the branch tips of the chicken lung were smaller, and the branch stalks were wider, only elongating and shaping into tubes later, possibly when surrounded by smooth muscle (Kim *et al*., 2015; Goodwin *et al*., 2022).

**Figure 1:**
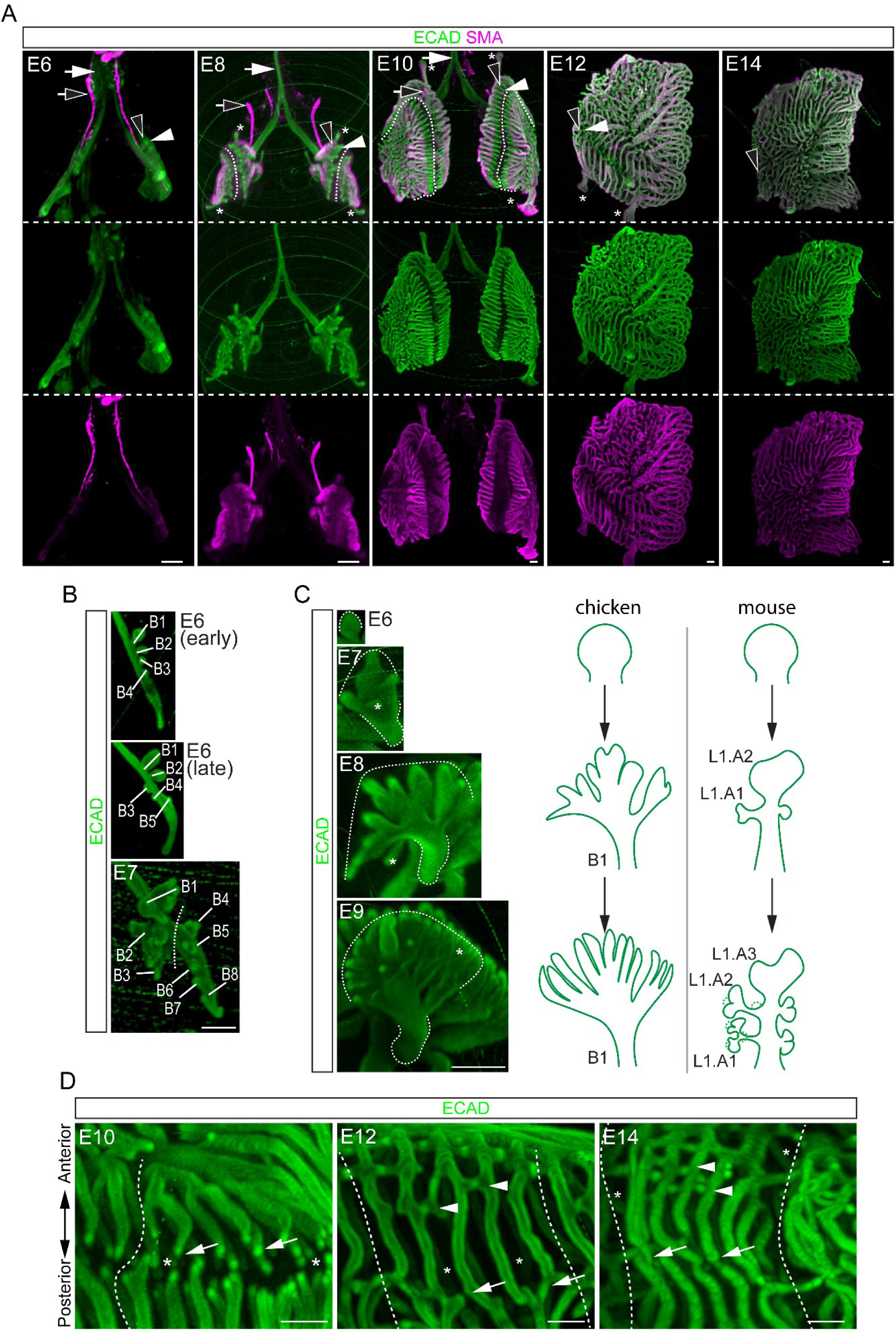
Multiscale time-course 3D imaging identifies proximal-short and distal-long fusions of hyper-elongated primary termini. (A) OPT images of wholemount immunostained chicken lungs during branching and fusion for the epithelium (ECAD, primary termini marked by filled arrowhead) and the airway (distal front of the same primary termini marked by open arrowhead) and vascular (open arrow) smooth muscle (SMA). Trachea and extrapulmonary airways are devoid of smooth muscle (filled arrow). Asterisk: air sac. Dash: gap between anterior and posterior groups of primary termini. Scale bars: 250 µm. (B) OPT images of chicken left lobes over time with labeled branches. The gap (dash) between anterior (B1-B3) and posterior (B4-B8) groups is obvious by E7. Scale bars: 250 µm. (C) OPT images of the chicken B1 branch outlined with dashes over time and schematics, in comparison with the bifurcating mouse lung. Asterisk: internal branches. Scale bars: 50 µm. (D) OPT images of chicken primary termini of the anterior group approaching and fusing with (arrow; distal-long fusion) those of the posterior group over time. Arrowhead: proximal-short fusion between tubes within the anterior group. Asterisk: internal parabronchi. Dash: sulcus. Scale bars: 250 µm.

By E7, a gap along the main bronchus widened to form an anterior group of side branches (B1, B2, and B3) and a posterior group (B4 through B8) (Fig. 1B). Arrays of parallel tubes from each group extended and bent at the front toward the opposite group, like two clapping hands with approaching fingertips (Fig. 1A, 1D). Within each group or clapping hand, parallel tubes also filled the palm region; these inter- and intra-groups of tubes possibly corresponded to paleopulmonic and neopulmonic regions, respectively, as reported for different avian species (SupFig. 1A) (Duncker, 1972). Regardless, unlike the mouse lung packed with perpendicular side branches and rosette-like orthogonal bifurcation (Metzger *et al*., 2008), the chicken lung had longer tubes with less interruption by side branches, possibly leaving space between parallel tubes for future alveoli.

The termini of the hyper-elongating branches, termed primary termini to distinguish them from subsequent neoform termini off the parabronchi, had different fates. A few of them, most noticeably that of the main bronchi and the most anterior branch (B1), ballooned after E8 and were easily severed during tissue dissection, corresponding to the storage air sacs introduced earlier (Fig. 1A asterisks). By E10, primary termini of the two opposing groups of parallel tubes aligned with remarkable synchrony across a ∼125 um gap with a limited standard deviation (30 um; 9 pairs of termini/lung and 3 lungs; Fig. 1A dashes; Table S1), reminiscent of microtubules attaching to invisible metaphase chromosomes – perhaps similarly dependent on organizing, attraction/repulsion, and/or checkpoint mechanisms (Maiato *et al*., 2017). These opposing primary termini fused between E12 and E14, which we named distal-long fusion to reflect their location (Fig. 1D arrows). Proximal to these, shorter branches budded off the whole length of the long branches, approached their counterparts off neighboring long branches, and also fused to bridge tubes, which we accordingly named proximal-short fusion (Fig. 1D arrowheads). Intragroup tubes of neopulmonic regions also fused both distally and proximally (SupFig. 1B). Thus, distal-long fusion converts blind-ended hyper-elongating branches into continuous long parabronchi that are short-circuited by proximal-short fusion, presumably accelerating the airflow at the expense of turbulence and mixing.

### Neoform termini protrude through parabronchial smooth muscle for radial alveologenesis

Our whole-lung imaging of epithelial tubes provided a 3D reference for mapping non-epithelial structures and cellular specifications. Chicken trachea and extrapulmonary bronchi were devoid of smooth muscles (Fig. 1A arrows). Also unlike the mouse lung where airway smooth muscle cells immediately wrapped and possibly promoted nascent branch clefts (Kim *et al*., 2015), chicken primary termini at the lobe edge outpaced smooth muscle actin expression by ∼180 um at E10 and ∼130 um at E12 until distal-long fusion, allowing smooth muscle cells to catch up and cover the entire parabronchi (Fig. 1A arrowheads; Fig. 2A; 6-9 termini each from 3 lungs at each developmental stage). The race between primary termini and smooth muscle cells was closer for proximal-short fusion (∼35 um at E12), possibly due to simple geometry, but ended still with smooth muscle-covered parabronchi (Fig. 1A, 2A).

**Figure 2:**
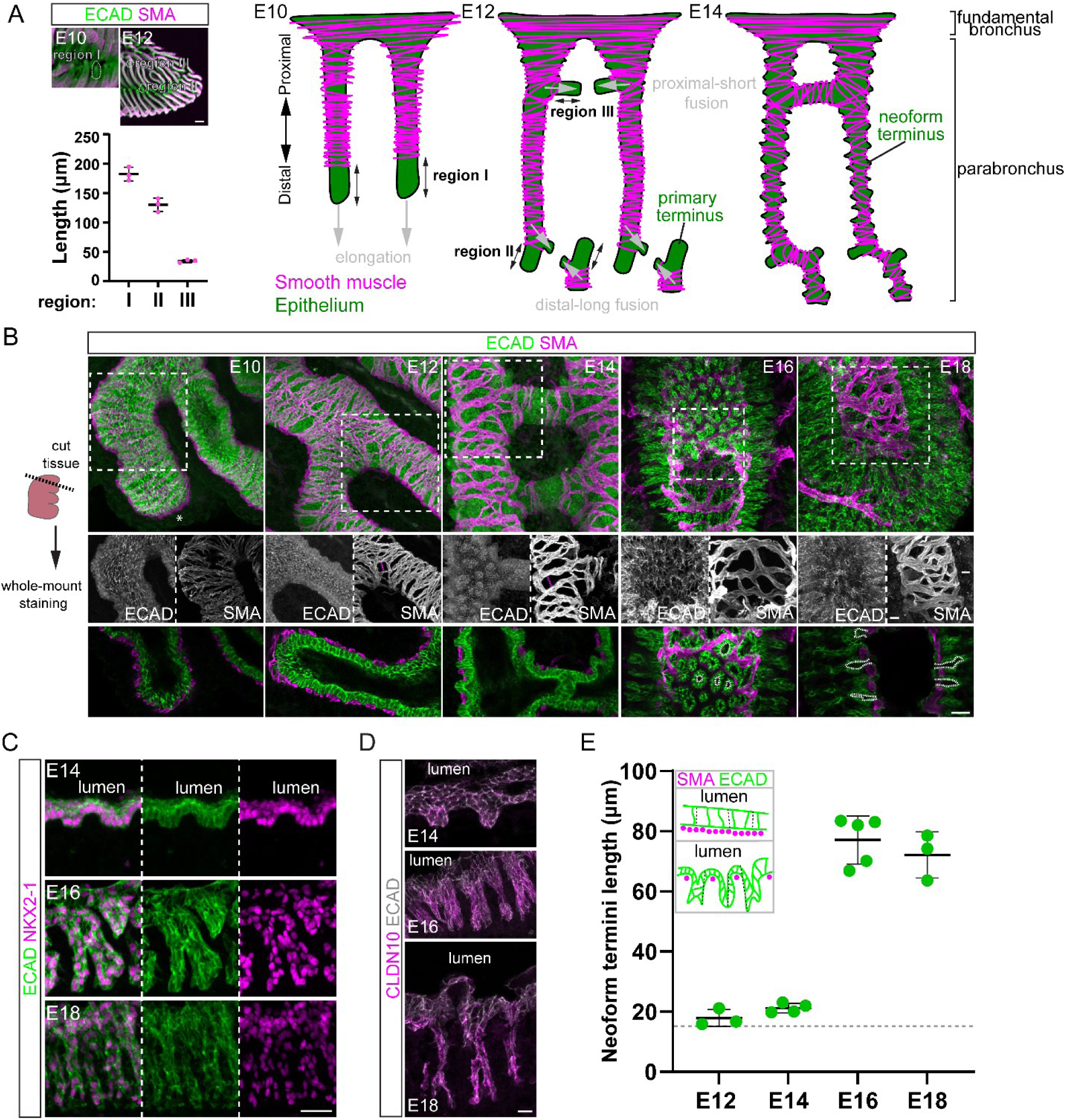
Neoform termini protrude through parabronchial smooth muscle for radial alveologenesis. (A) Diagram of chicken lung epithelium and smooth muscle over time. Black double arrow: length of the epithelium without smooth muscle coverage of indicated regions measured in the plot; each symbol represents one lung with at least 6 branches per lung. E10 and E12 OPT images illustrate the 3 measured regions. Scale bars: 250 µm (B) Confocal projection (top two rows) and optical section (bottom row) images of chicken lung strips cut along the first costal sulcus (diagram), showing the epithelium (ECAD) and smooth muscle (SMA). Gaps between smooth muscle fibers (magenta dash in middle row) widen from E12, coinciding with epithelial protrusion. White dash: lumen of neoform termini. Asterisk: epithelium out of imaging plane. Scale bars: 25 µm. (C) Confocal images of chicken lung sections showing neoform termini led by individual NKX2-1+ epithelial (ECAD+) cells. Scale bars: 25 µm. (D) Confocal images of chicken lung sections immunostained for an epithelial junction marker (ECAD) and a more diffuse epithelial marker (CLDN10), showing radial extension of neoform termini. Scale bars: 10 µm. (E) Quantification of radial outgrowth of neoform termini. Inset: schematic of measured distance (dash). Horizonal dash denotes the baseline thickness of epithelial cells at E10 prior to radial outgrowth. Each symbol represents one lung with 15-20 measurement points.

To achieve cellular resolution and image post-fusion lungs that were too complex for OPT, we switched to whole-mount confocal imaging of tissue strips cut along the first cranial costal sulci that could be reliably identified (Fig. 2B) (Palmer and Nelson, 2020). At E10, the columnar parabronchial epithelium was confined within tightly weaved SMA+ smooth muscle fibers, which dispersed subsequently to form increasingly larger gaps, either from active cell rearrangement or passive tube elongation (Fig. 2B).

Through these gaps, the epithelium protruded radially and elongated to form what we called neoform termini to reflect their blind-ended nature and origin from non-terminal tubes (Fig. 2A, 2B). From snapshots of the initiation, progression, and resolution phases of primary termini fusion with increasing smooth muscle coverage, neoform termini formed outside the fusion zone (SupFig. 2). The narrow lumen inside neoform termini would aerate the future acinus, infundalubus, and alveolus radially from the unidirectional parabronchus. Unlike primary termini and those of the mouse lung, neoform termini had an irregular leading edge consistent with migration or invasion of a few cells (Fig. 2C). The radial outgrowth of neoform termini was better visualized by immunostaining for an epithelial claudin, CLDN10 (La Charité-Harbec *et al*., 2022), that localized to cell junctions as well as intracellularly (Fig. 2D, 2E). Therefore, the chicken lung undergoes radial alveologenesis via protrusion of neoform termini in regions devoid of smooth muscle.

### Branch stalks halt proximalization to form parabronchi and neoform termini

Generation of new distal ends via outgrowth of neoform termini from otherwise proximal parabronchi disrupted the tight coupling between branching and proximalization of the mammalian lung. Throughout most of the embryonic mouse lung development, branch stalks left behind by SOX9+ branch tip progenitors expressed SOX2, a marker and promoter of airway differentiation (Alanis *et al*., 2014), such that no gap existed between SOX9 and SOX2 expression domains (Fig. 3A, 3B, SupFig. 3A). Only after E16.5, even though SOX9 progenitors continued to branch, no more branch stalks turned on SOX2; a gap between SOX9 and SOX2 formed and widened, marking the future alveolar ducts that consisted of alveolar rather than airway cells (SupFig. 3B) (Alanis *et al*., 2014).

**Figure 3:**
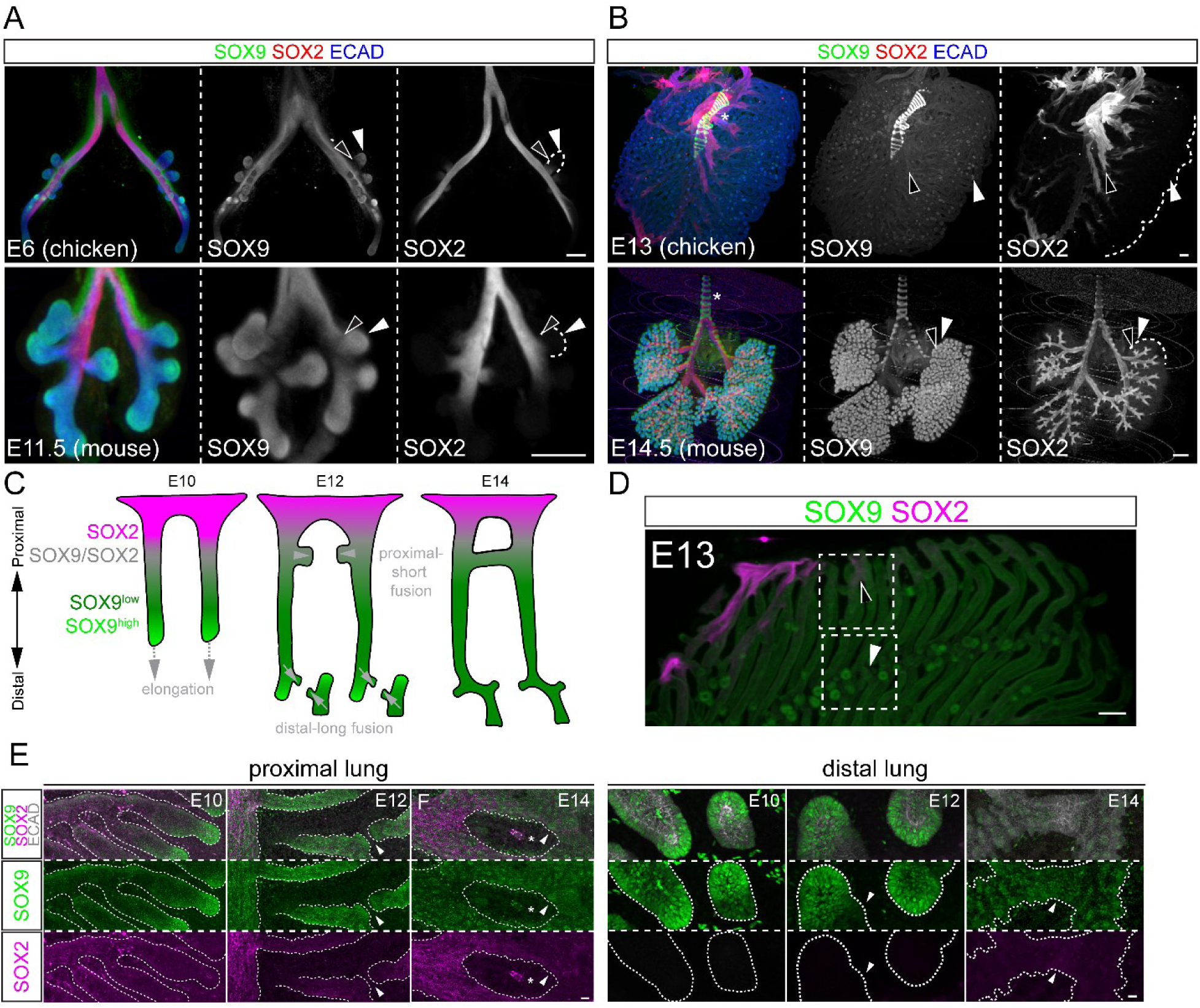
Branch stalks halt proximalization to form parabronchi and neoform termini. (A-B) OPT images of immunostained chicken and mouse lungs at comparable developmental stages for the epithelium (ECAD; dash) and its distal (SOX9) and proximal (SOX2) markers, as well as their respective fronts (filled and open arrowheads). Asterisk: SOX9^+^ cartilage rings. Scale bars: 250 µm. (C) Schematic to illustrate SOX2 and SOX9 expression during proximal-short fusion and distal-long fusion. (D) Representative OPT image of the boxed proximal and distal regions for Fig. 3E. Open arrowhead: proximal-short fusion. Filled arrowhead: distal-long fusion. Scale bars: 250 µm (E) Confocal images of immunostained chicken lungs mounted to view the proximal (top) and distal (bottom) regions of the parabronchi. Proximal-short, but not distal-long, fusion (arrowhead) is adjacent to SOX2 expression. SOX9^high^ primary termini are followed by SOX9^low^ branch stalks. Scale bars: 25 µm proximal, 15 µm distal.

By comparison, E6 chicken lungs with comparable branch complexity to E11.5 mouse lungs, also had SOX2+ branch stalks immediately behind SOX9+ branch tips, suggesting a conserved proximalization mechanism at this early stage of lung development (Fig. 3A). However, by E13, chicken branch stalks no longer activated SOX2 and a large gap formed behind the SOX9^high^ elongating branch tips, in contrast to the juxtaposition of SOX9 and SOX2 in E14.5 mouse lungs (Fig. 3B arrows). Branch stalks in this gap expressed a lower level of SOX9 and were the future parabronchi that would give rise to alveoli via neoform termini – possibly a reason for not expressing the airway-promoting SOX2 and a more efficient design than turning off SOX2 later, as reported for the snake lung (van Soldt *et al*., 2024) (Fig. 3B, 3C, 3D). At a cellular resolution and on cross-sections, SOX9^low^ branch stalks grew neoform termini, often led by SOX9-expressing cells, before downregulating SOX9 and beginning alveolar differentiation, a switch from progenitor to differentiation programs also observed in the mouse lung (Chang *et al*., 2013; Rockich *et al*., 2013) (SupFig. 3C).

SOX2 expression molecularly corroborated our structural classification of proximal-short and distal-long fusions (Fig. 3C). Proximal-short fusions involved branches close to the SOX2 expression domain such that the fusing epithelium contained SOX2+ cells, whereas distal-long fusions had no SOX2 expression (Fig. 3D, 3E). Consistent with prior reports (Palmer and Nelson, 2020), approaching SOX9^high^ tips narrowly missed each other and fused laterally, suggesting an uncoupling between elongation and fusion as well as a signal possibly from fusion to stop growth and eliminate branch tips (Fig. 3E).

Taken together, proximalization of branch stalks, as measured by SOX2 expression, proceeds during early chicken lung development to form fundamental bronchi (Fedde, 1998), but halts mid-development to form SOX2-SOX9^low^ parabronchi, from which neoform termini grow radially to form alveoli.

### Primary vascular plexus matures into macrovasculature, off which secondary vascular plexus extends to form capillaries

As epithelial fusion eliminated primary termini and radial outgrowth reestablished neoform termini – redefining the proximal-distal axis, the vasculature must adapt to ensure juxtaposition with the epithelium at the air capillaries. Similar to the mouse lung (Lazarus *et al*., 2011), primary termini of the E6 chicken lung were encased by a vascular plexus marked by ERG and CLDN5 (Fig. 4A). This primary plexus was connected with proximal vascular tubes that paralleled the airways and were arterial (marked by SOX17) and venous macrovasculature based on location and smooth muscle coverage (Fig. 1A, 4A). Branching and elongation of primary termini were accompanied by the growth of the primary vascular plexus so that by E10, approaching tips of the future parabronchi were capped by a merged plexus whereas their proximal flanks were paralleled by remodeled vascular tubes, consistent with a continuous conversion of plexi into tubes to establish a vascular tree (Fig. 4B).

**Figure 4:**
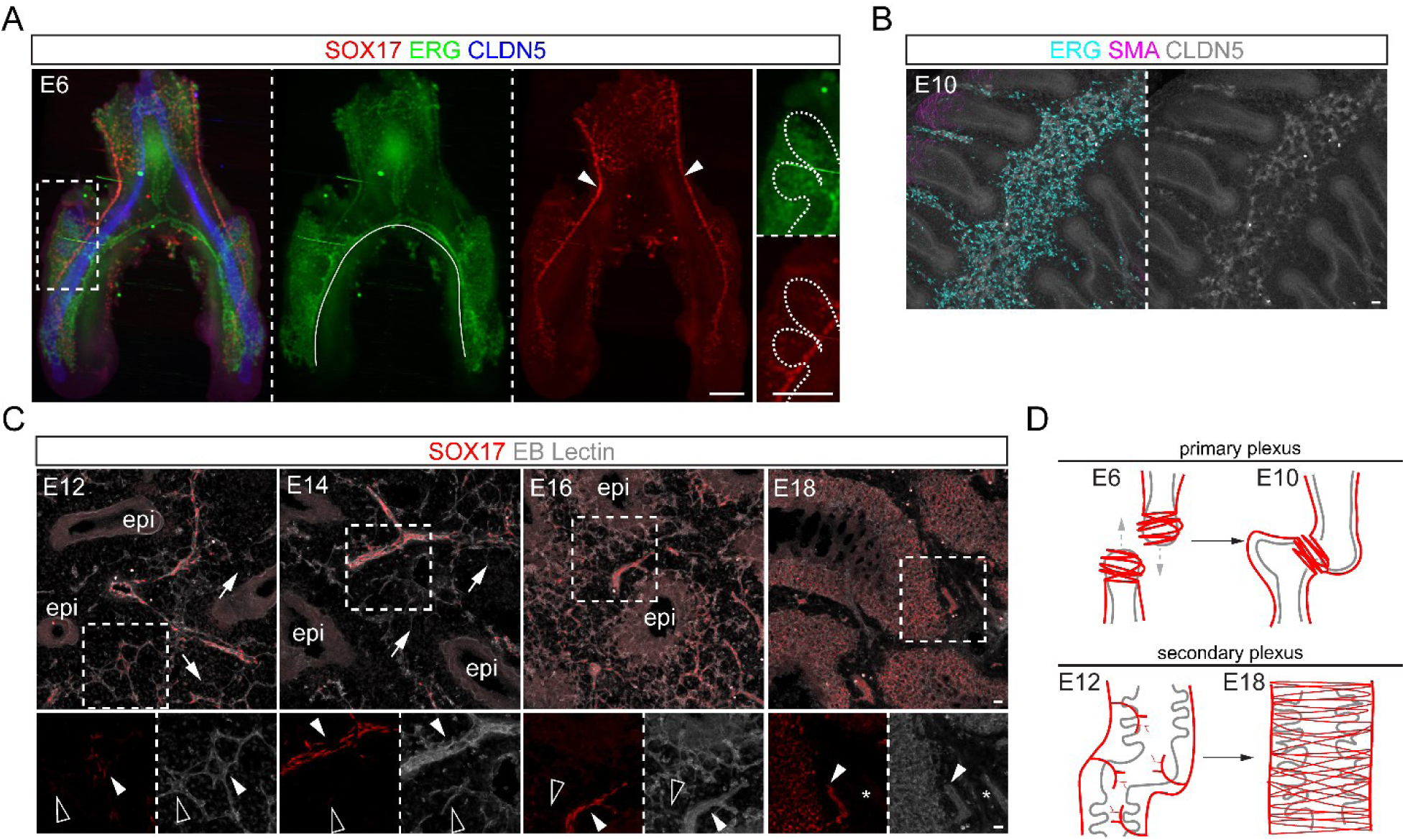
Primary vascular plexus matures into macrovasculature, off which secondary vascular plexus extends to form capillaries. (A) OPT images of an E6 chicken lung immunostained for the vasculature (ERG), arterial macrovasculature (SOX17; arrowhead), and epithelium (CLDN5, low in OPT for vasculature). Solid line: venous macrovasculature based on morphology and location. Dash: traced epithelium in the boxed region. Scale bars: 250 µm. (B) Confocal images of a wholemount immunostained E10 chicken lung showing that the primary plexus (ERG+; CLDN5+, visible on confocal) caps individual branch tips (CLDN5+) and merges with those of adjacent and opposing branch tips. Branch stalks (CLDN5+) surrounded by smooth muscle (SMA) alternate with remodeled tubular vessels (ERG+; CLDN5+). Scale bars: 20 µm. (C) Confocal images of chicken lung sections showing the emergence of the secondary plexus (EB lectin+ SOX17-; open arrowhead) around the remodeled primary plexus (EB lectin+ SOX17+; filled arrowhead). At E18, the secondary plexus interdigitating with neoform termini becomes SOX17+. Asterisk: venous (SOX17-) macrovasculature. Epi: epithelium. Arrow: sprout-like structures. Scale bars: 25 µm. (D) A diagram of primary and secondary plexi and their relationship to the macrovasculature and the epithelium (grey).

At E12, while the plexus-to-tube remodeling lingered around the pre-fusion tips (SupFig. 4A), the remodeled proximal vessels, which expressed SOX17 and situated halfway between epithelial tubes on sections, began to grow a secondary plexus (EB lectin+) that over time interdigitated with the outgrowing neoform termini (Fig. 4C, 4D). Unlike the primary plexus, SOX17 expression extended to the secondary plexus (Fig. 4C). Additional endothelial markers ERG and CAV1 also suggested the origination of the secondary plexus from remodeled vessels and also captured sprout-like intermediates, supporting the notion that sprouting angiogenesis seeded the secondary plexus, as proposed by classical electron microscopy studies (West, Bamford and Jones, 1977; Makanya *et al*., 2007; Makanya and Djonov, 2009; West *et al*., 2010) (SupFig. 4B, 4C). Therefore, sprouting a secondary plexus from the proximal macrovasculature topologically matches the out-pocketing of neoform termini from the proximal parabronchi (Fig. 4C), unlike the mammalian lung where the primary plexus occupies the same distal region as branch tips that eventually transform into alveoli.

### ScRNA-seq defines epithelial progenitors and a third, chicken-specific alveolar cell type

To gain unbiased molecular insights into conserved and species-unique processes of lung development, we leveraged single-cell genomics technology that is applicable to any species of a sequenced genome and allows cross-species comparison. E13 and E20 chicken lungs, amid tubulogenesis and alveolar differentiation, respectively, were profiled and compared to published mouse lung scRNA-seq data from equivalent stages, E14.5 and E18.5 (Gerner-Mauro, Akiyama and Chen, 2020; Vila Ellis *et al*., 2020). Realizing that chicken erythrocytes were large, nucleated, and resistant to hypotonic lysis, we used FACS to exclude them by size. We also enriched for epithelial cells underrepresented from tissue dissociation by immunostaining for an extracellular epitope of E-Cadherin (ECAD) (SupFig. 5A). Furthermore, chicken erythrocytes were PTPRC (CD45) negative such that CD45 depletion commonly used for mammalian lungs was ineffective in removing them; the resulting scRNA-seq data also verified that our E-Cadherin enrichment sorting strategy did not miss major epithelial populations (SupFig. 5B).

At E13, 3628 computationally purified epithelial cells (*CDH1*+*CDH5*-*COL3A1*-*PTPRC*-; additional low-quality cells removed after subsetting and reclustering) formed 4 clusters marked by *SOX9*, *TOP2A*, *FOXJ1*, and *KRT14* (Fig. 5A, 5B, 5C, 5D; Table S2). Correlating with cell proportions in immunostaining and comparing to published mouse E14.5 scRNA-seq data (Gerner-Mauro, Akiyama and Chen, 2020), the largest *SOX9*+ cluster corresponded to cells of primary termini and parabronchi, adjacent to and thus related to the proliferative cluster (Fig. 5B). A sub-cluster within had higher *SOX9*, *FGFR2*, *SFTPC*, and more transcripts in general, suggesting a more active state potentially in cells undergoing morphogenesis such as elongation, fusion, and radial protrusion (Fig. 5D, SupFig. 5C). Intriguingly, neuronal guidance molecules, *ROBO2* and *SLIT3*, were present in chicken, but not mouse, *SOX9*+ cells, raising speculation of chemoattraction or repulsion operating on multicellular structures to control termini growth and fusion (Fig. 5D, 5E). This conserved and unique gene expression was captured via direct quantitative comparison of name-based orthologs in chicken and mouse SOX9 cells (Fig. 5E). The *SOX2*+ cluster was expectedly much smaller than its mouse counterpart (Gerner-Mauro, Akiyama and Chen, 2020) given the halted proximalization and consisted mainly of ciliated cells (Fig. 5D, SupFig. 5E). The *KRT14*+ cluster was considered the precursor to a new cell type examined in detail below.

**Figure 5:**
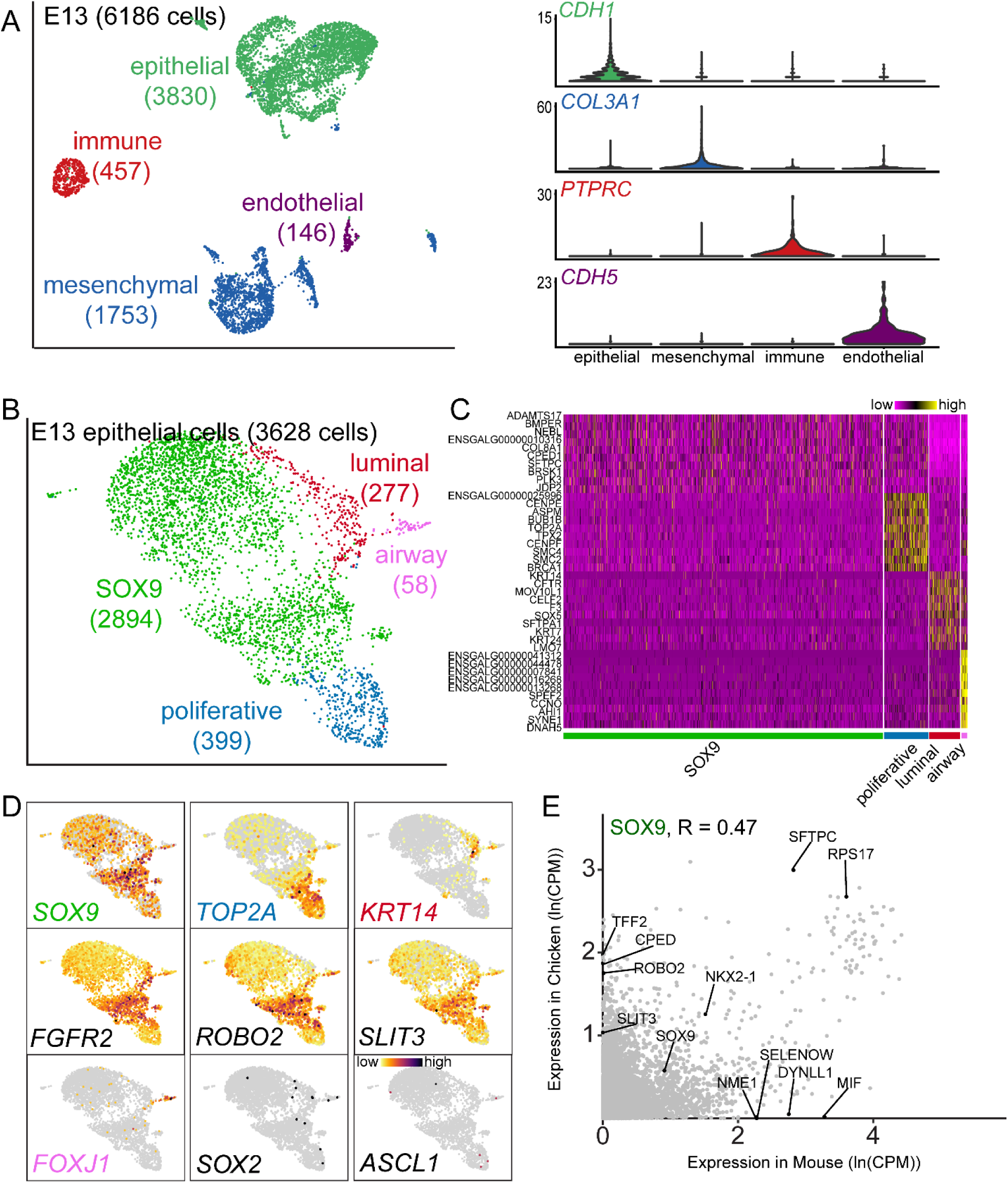
ScRNA-seq profiling of E13 chicken lung epithelial progenitors identifies species shared and unique features. (A) UMAP (left) and violin (right) plots of ECAD-enriched E13 chicken lung cells with the cell number of each lineage in parentheses. (B) UMAP plot of computationally purified E13 chicken epithelial cells (*CDH1*+) with the cell number of each cell type in parentheses. (C) Heatmap of top 10 genes of each epithelial cell type. (D) Feature plots of color-coded, cell-type-specific genes and chicken-specific axonal guidance genes (*ROBO2* and *SLIT3)*. *SOX2* (airway marker) and *ASCL1* (neuroendocrine marker) are too low to be informative. (E) Scatterplot comparison of expression levels of common genes in E13 chicken and E14.5 mouse SOX9 cells. CPM: count per million. R: Pearson correlation coefficient.

At E20, one day before hatching when alveolar differentiation was sufficiently complete for breathing (Dawes, 1976; Akiyama *et al*., 1999), 1989 computationally purified epithelial cells formed 3 clusters – one more than the mouse lung at E18.5 (Vila Ellis *et al*., 2020), one day before birth, consisting of alveolar type 1 (AT1) and type 2 (AT2) cells responsible for gas exchange and surfactant production, respectively (Liem, 1988) (Fig. 6A, 6B, 6C, SupFig. 5E, Table S2). Conserved, specific expression of *VEGFA*, *IGFBP7*, *AQP1*, *LAMP3*, *SFTPC*, and *LGI3* supported the most parsimonious naming of 2 clusters as chicken AT1 and AT2 cells, the latter of which included a small proliferative *TOP2A*+ subcluster (Fig. 6C, 6D). Systematic comparison of name-based orthologs in AT1 and AT2 cells across species identified shared genes, such as *VEGFA* in AT1 and *SFTPA1* in AT2 cells, as well as expected housekeeping genes and pan-epithelial genes (*CDH1* and *NKX2-1*) (Fig. 6E). Among alveolar cell-type-specific genes, *LY6E* gained and *AQP5* lost AT1-specificity in chicken; *FABP3* gained and *DLK1* lost AT2-specificity in chicken (Fig. 6E). Such shift in cell-type-specificity likely reflected promoter/enhancer evolution and had been categorized as gain/loss, expansion/contraction, or switch when factoring in additional cell types in mouse versus human lungs (Travaglini *et al*., 2020).

**Figure 6:**
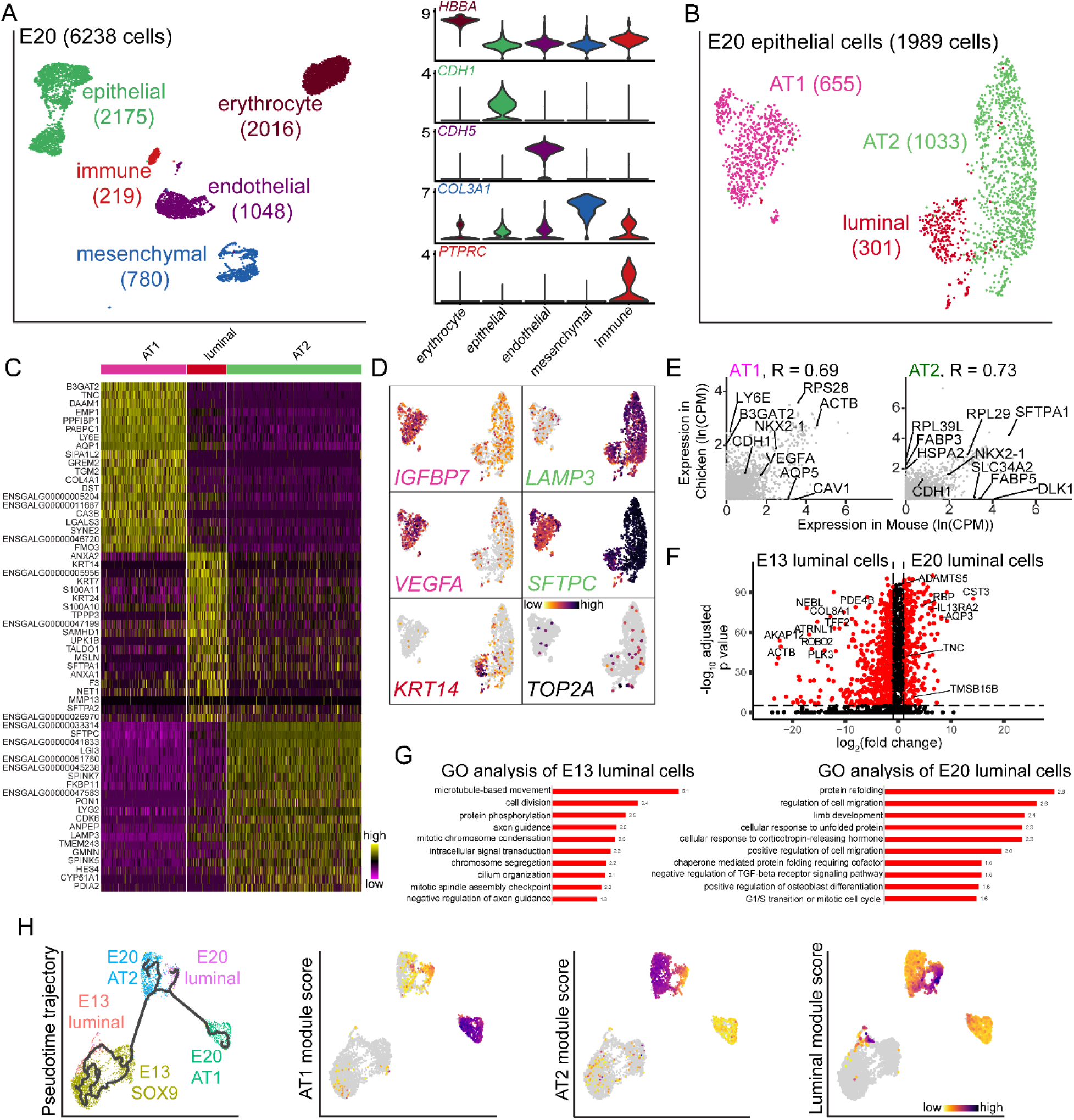
ScRNA-seq identifies a third, chicken-specific alveolar cell type. (A) UMAP (left) and violin (right) plots of ECAD-enriched E20 chicken lung cells with the cell number of each lineage in parentheses. (B) UMAP plot of computationally purified E20 chicken epithelial cells (*CDH1*+) with the cell number of each cell type in parentheses. (C) Heatmap of top 20 genes of each epithelial cell type. (D) Featureplots of color-coded cell-type-specific genes for AT1, AT2, and luminal cells, as well as proliferative cells (*TOP2A*). (E) Scatterplot comparison of expression levels of common genes in E20 chicken and E18.5 mouse AT1 and AT2 cells. CPM: count per million. R: Pearson correlation coefficient. (F, G) Volcano plot comparison and GO analysis of E13 and E20 luminal cells. (H) Monocle pseudotime analysis of aggregated E13 and E20 lung epithelial cells, excluding airway cells at E13, showing a trajectory from E13 SOX9 cells to each of the E20 alveolar cell types. Overlaid on the same UMAP, module scores of top 20 cell-type-specific genes in (C) show luminal, but not AT1 nor AT2, cell differentiation at E13.

Although the third, chicken-specific cluster was near the AT2 cluster and also had, albeit at a lower variable level, *LAMP3* and *SFTPC*, it uniquely expressed *KRT14*, the same marker for the unaccounted-for cluster at E13. This *KRT14*+ cluster was considered distinct from AT2 cells and named luminal cells, as supported by their tissue location examined below (Fig. 5, Fig. 6C, 6D, SupFig. 5D). Comparison of E13 and E20 luminal cells revealed a maturation process with downregulation of microtubule and axon guidance genes such as *ACTB* and *ROBO2* and upregulation of cell migration genes such as *TMSB15B* (Fig. 6F, 6G). Only the gene signature of luminal cells, not that of AT1 or AT2 cells, was apparent at E13 (Fig. 6H). Trajectory analysis of *SOX9*+, luminal, and AT1 and AT2 cells supported *SOX9+* cells as progenitors, reminiscent of the SOX9+ bipotential progenitors of the mouse lung, although definitive lineage tracing experiments in the chicken lung were not performed (Fig. 6H).

### Alveolar type 1, type 2, and luminal cells occupy concentric zones around parabronchi

To map AT1, AT2, and luminal cells amid dynamic radial alveologenesis, we performed time-course RNAscope for their respective markers: *VEGFA*, *LAMP3*, and *KRT14*. At E14 during initial protrusion of neoform termini, the markers had a sporadic and intermixed distribution, with at least some in the same cell (Fig. 7A). Over time, they segregated into concentric zones of *VEGFA*, *LAMP3*, and *KRT14* expression surrounding the lumen in a centripetal order (Fig. 7A). As a result, AT1 cells preferentially coated air capillaries, and AT2 cells infundibula and atria; *KRT14*+ cells lined the parabronchial lumen, thereby named luminal cells (Fig. 7B). This cell-type-specific zonation corroborated the presence of the third, chicken-specific alveolar cell type identified by scRNA-seq and, in hindsight, might reflect an optimal cell distribution from radial alveologenesis where distalmost AT1 cells mediated gas exchange, surfactant-producing AT2 cells conditioned air sac entryways, and luminal cells, excluded from outgrowth, lined the parabronchial canals. Luminal cells were not a developmental intermediate because they still lined the parabronchial lumen in the adult chicken lung, albeit in a broader domain including the atria (Fig. 7C). In the mouse lung, however, KRT14 was detected in basal cells in proximal bronchi and mesothelial cells on the tissue surface, but not in the alveolar region (SupFig. 7).

**Figure 7:**
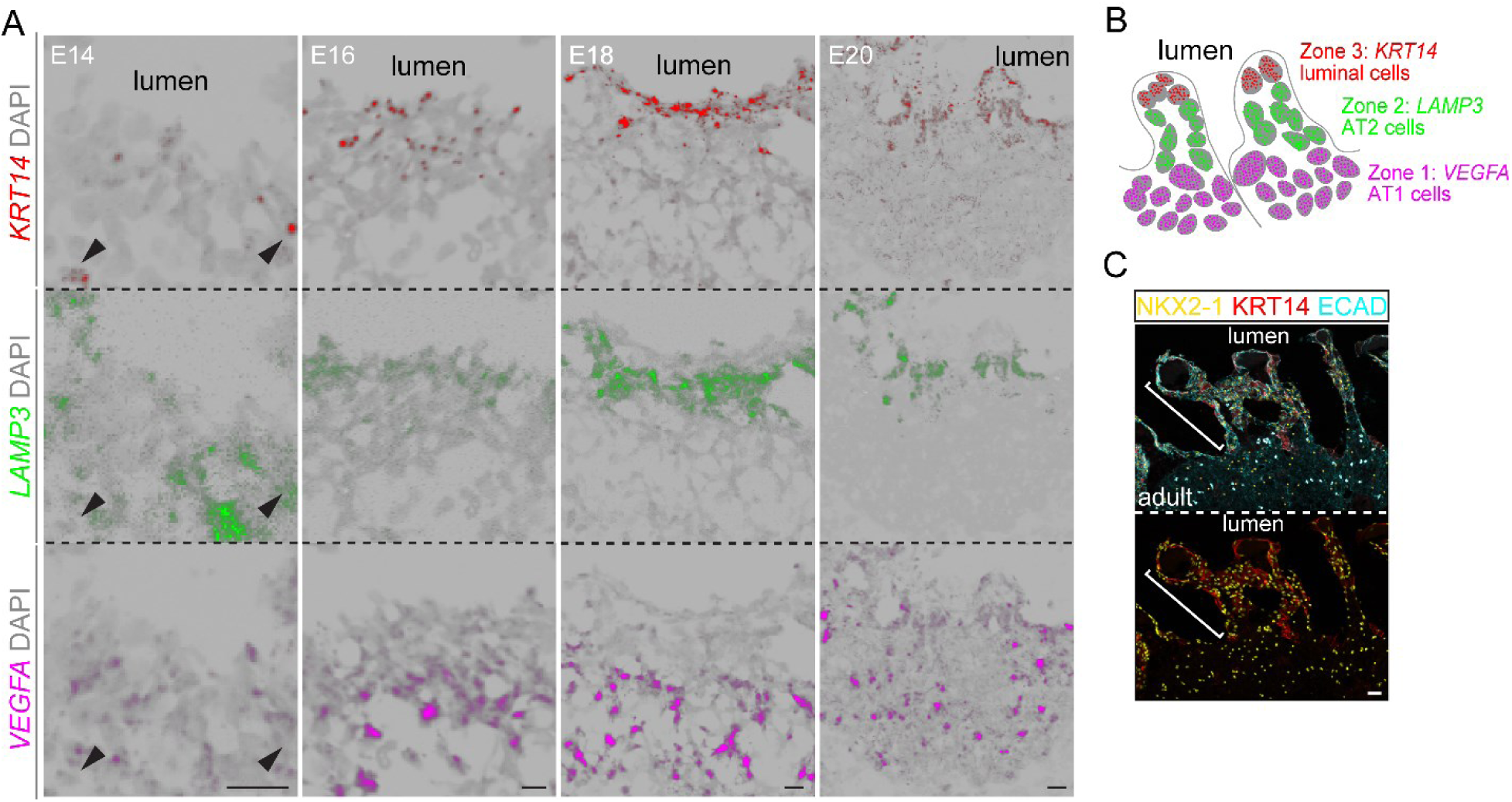
AT1, AT2, and luminal cells occupy concentric zones around parabronchi. (A-B) Time-course RNAscope images for AT1 (*VEGFA*), AT2 (*LAMP3*; lower in luminal cells), and luminal cells (*KRT14*), showing the formation of cell type zonation (diagram in B). Arrowhead: marker co-expression in cells of early post-fusion lungs. Scale bars: 10 µm. C) Confocal images of an adult chicken lung section immunostained for lung epithelial nuclei (NKX2-1) and junctions (ECAD), showing luminal cells (KRT14) boarding the lumen including the atria (bracket). Scale bars: 25 µm.

To extend our localization analysis beyond individual markers, we leveraged Stereo-seq to measure the whole transcriptome at a resolution of 0.5 um on an E20 chicken lung section (Fig. 8A). Considering the limitation of square bins in capturing the diverse, interdigitated cell types in the lung, we assigned 2.73 billion sequencing reads to 315,012 segmented cell bins with a mean measurement point of 118 and a mean gene number of 174. This resulted in a structured UMAP albeit without discrete clusters, likely due to low gene coverage and imprecision in cell segmentation (Fig. 8A).

**Figure 8:**
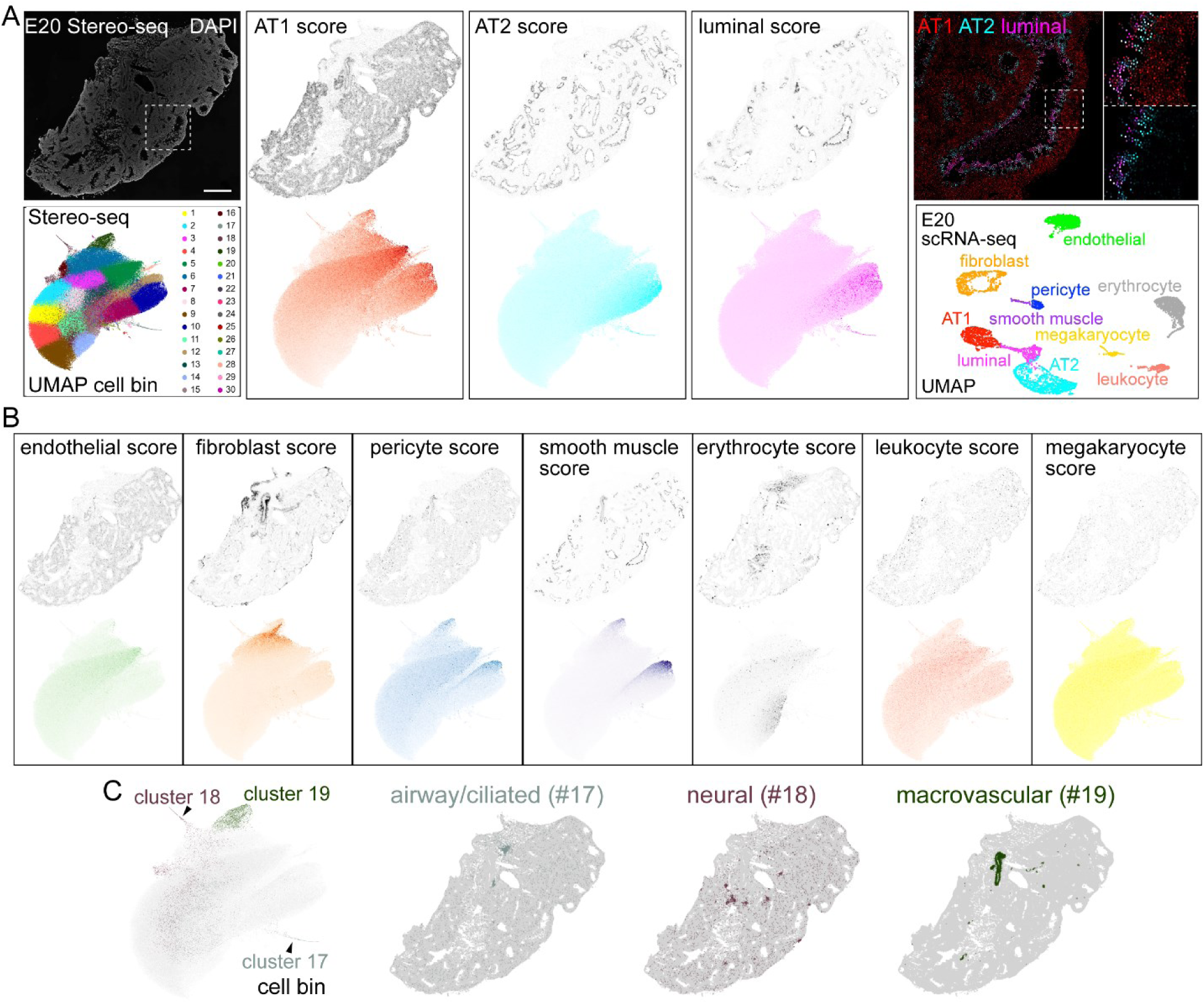
Stereo-seq spatial transcriptomics localization of chicken lung cell types. (B) Left column: E20 Stereo-seq section nuclei (DAPI) image and cell bin UMAP with its default 30 clusters. Right columns: tissue (top) and UMAP (bottom) display of module scores of AT1, AT2, and luminal cells based on top 20 genes of E20 scRNA-seq. Boxed regions are sequentially enlarged to visualize cell type zonation, with module scores color-coded as fluorescence intensity. Scale bars: 1 mm. (C) Tissue (top) and UMAP (bottom; color-matched with scRNA-seq) display of top 20-gene module scores of remaining cell types captured by scRNA-seq. (D) Small but distinct clusters of cell bin UMAP (left) correspond to low-abundance cell types/niches with characteristic tissue distributions (right) missed by scRNA-seq.

Accordingly, instead of individual genes, we generated for each cell bin gene module scores using top 20 genes of each cell type captured by scRNA-seq (Fig. 8A, 8B, Table S3). Consistent with our RNAscope data (Fig. 7A), cell bins with high AT1, AT2, and luminal gene scores occupied concentric zones around the parabronchial lumen (Fig. 8A). Similarly, we localized endothelial, mesenchymal, and immune cell types, corresponding to distinct regions of the cell bin UMAP (Fig. 8B). Stereo-seq cell bin clusters also included cells/niches of low abundance and thus not captured by scRNA-seq; their tissue distribution and marker genes were consistent with airway/ciliated (cluster 17), neuronal (cluster 18), and macrovascular (cluster 19) structures (Fig. 8C; Table S3).

## DISCUSSION

In this study, we have delineated the morphogenesis of air passages throughout embryonic chicken lung development, including coordination of epithelial, smooth muscle, and vascular structures (Fig. 1, Fig. 2, Fig. 4). Hyperelongating primary termini undergo long-distal and short-proximal fusion and leave behind SOX9^low^ SOX2-, muscle-wrapped branch stalks as future parabronchi, from which neoform termini out-pocket radially. The primary vascular plexus encases each primary terminus and remodels at the midpoint between parabronchi into SOX17+ macrovasculature. The secondary vascular plexus seemingly sprouts from this macrovasculature and interfaces with neoform termini to form alveoli (Fig. 4C, 4D). We have also profiled chicken lung epithelial cells at key developmental stages and identified equivalent cell types of the mouse lung as well as a third, chicken-specific alveolar cell type, the luminal cell, that occupies together with AT1 and AT2 cells concentric zones, possibly patterned by radial alveologenesis (Fig. 6, Fig. 7, Fig. 8). Our study sheds light on the morphogenic, molecular, and cellular processes supporting the unidirectional flow in the chicken lung and illustrates the use of 3D imaging and single-cell genomics in comparative evolutionary biology of species-unique features, potentially even in non-model organisms.

The contrasting difference between the tree-like E14.5 mouse lung and racetrack-like E13 chicken lung (Fig. 3B) can be traced to discrete behaviors of their branch tips. Even at an initial stage when comparable lateral branches form in the E11.5 mouse lung and E6 chicken lung, the chicken main bronchi are much longer (Fig. 3A).

This tendency to elongate rather than bifurcate continues until fusion of primary termini. Such species-specific adaptive change in the relative rates of elongation and bifurcation is operative across branching organs of the same species, such as the hyper-bifurcating kidney and hyper-elongating mammary glands (Richards *et al*., 2004; Menshykau *et al*., 2019). On the molecular level, FGF10 signaling regulates branching of all three organs (Bellusci *et al*., 1997; Michos *et al*., 2010; Zhang *et al*., 2014), and must be tuned spatiotemporally or modulated by other signals such as GDNF in the kidney (Costantini, 2010). Elongation is much less understood and depends on convergent extension in the kidney collecting ducts that remarkably resemble the chicken parabronchi. Airway smooth muscle surrounding the parabronchi might also provide a circumferential constraint to favor longitudinal cell alignment. Little is known about fusion of multicellular tubes including the contribution and control of apoptosis in resolving aggregated cells (Ray and Niswander, 2012) and the absence of fusion despite tight packing of branch tips in the mouse lung. Functional studies of chicken-specific molecules from our scRNA-seq profiling, such as ROBO and SLIT, are expected to be informative (Fig. 5D).

The bimodal expression of distal-SOX9 and proximal-SOX2 in the lung is typically called proximal-distal patterning, a term borrowed from classical *Drosophila* embryo patterning and implying a morphogen gradient (Karlsson, 1980; Cifuentes and García-Bellido, 1997). However, rather than patterning a preformed embryo, branch tips continue to expand distally and leave behind branch stalks that switch from SOX9 to SOX2 expression. The structural dynamics and expression switch absent in traditional patterning support proximal specification – activation of the proximal marker SOX2 – as the basis of SOX9-SOX2 expression in the lung. Although both mouse and chicken branch tips express SOX9 and grow the epithelial tree, proximal specification is halted in the mid-stage chicken lung but not the mouse lung, forming SOX9^low^ parabronchi such that radial alveologenesis does not originate from SOX2+ airways (Fig. 3D). By comparison, the snake lung seems to continue proximal specification but subsequently downregulates SOX2 in preparation for its radial alveologenesis (van Soldt *et al*., 2024); SOX2 is even transiently expressed in early branch tips of the human lung (Danopoulos *et al*., 2018). Mechanistically, *Sox9* or *Sox2* deletion in the mouse lung does not affect the other’s expression domain, consistent with an upstream regulator, such as Fgf signaling, that maintains SOX9 expression and suppresses proximal specification (Gontan *et al*., 2008; Alanis *et al*., 2014). Accordingly, expanded Fgf signaling beyond branch tips or the absence of a proximalization signal might enable the halted proximal specification in the chicken lung. Besides candidate gene approaches, future systematic dissection of enhancers of *Sox9* and *Sox2* and associated transcription factors could be informative.

The mouse gas exchange surface is covered with two intermixed alveolar cell types whose differentiation from a common pool of SOX9 progenitors is coupled with dilation of branch tips under the combinatorial transcriptional control of NKX2-1, YAP/TAZ/TEAD, and CEBPA (Little *et al*., 2021; Hassan and Chen, 2024). The chicken gas exchange surface forms via radial growth of the parabronchial epithelium, resulting in zoned, rather than intermixed, alveolar cell types (Fig. 7A, 7B, Fig. 8A). AT1 cells occupy the outermost zone to approach/attract the sprouting secondary vascular plexus possibly by producing the angiogenic factor VEGFA, supporting the notion that a second defining characteristic of AT1 cells besides their ultrathin morphology for gas diffusion is to signal toward the vasculature to establish the blood-gas interface. More speculatively, the secondary plexus approaching AT1 cells in the chicken lung is reminiscent of Cap2 endothelial cell differentiating in contact with AT1 cells in the mouse lung (Vila Ellis *et al*., 2020). AT2 cells occupy the entryway of individual alveoli, possibly sufficient to supply each comparatively small and isolated alveolus with surfactants without competing with AT1 cells for capillaries. If the AT1-AT2 intermixing in the mouse lung results from stochastic activation of YAP/TAZ and AT1 cell differentiation by mechanical stretching from alveolar expansion (Little *et al*., 2021; Penkala *et al*., 2021), alternative niche-dependent mechanisms could drive zone-specific AT1 and AT2 differentiation in the chicken lung.

Most notably, chicken-specific luminal cells occupy the innermost zone along the parabronchial lumen (Fig. 7, 8). Structurally, luminal cells are left behind by the outgrowth of neoform termini and are expected to experience sheer force from the unidirectional airflow, possibly as a result of expressing unique mechano-keratins including KRT14 as keratinocytes of the skin do (Ramms *et al*., 2013; Guo *et al*., 2020). Molecularly, KRT14 is a marker of mammalian basal cells, which function as dedicated epithelial stem cells (Ievlev *et al*., 2023). However, other basal cell markers including P63 are absent in chicken luminal cells, leaving their function an open question and calling for lineage tracing tools and experiments during development and recovery from injuries such as avian flu. Intriguingly, respiratory bronchioles and alveolar ducts in the mammalian lung also undergo out-pocketing (Yang *et al*., 2016; Basil *et al*., 2022), suggesting the possible existence of luminal cells or equivalent proximal-distal intermediates such as bronchioalveolar stem cells and respiratory airway secretory cells (Kim *et al*., 2005; Basil *et al*., 2022).

## MATERIALS AND METHODS

### Chickens (Gallus Gallus)

Single Comb White Leghorn fertilized eggs were supplied by the Texas A&M University Poultry Science department. The day of shipment was marked as embryonic day 0 (E0). Animals were incubated at 38°C for up to 20 days gestation. Adult chicken lungs were donated by The Groves Family Farm in Alvin, TX. All animal experiments were overseen and approved by the institutional animal care and use committee at The University of Texas MD Anderson Cancer Center.

### Antibodies

The following antibodies were used for the chicken data: rabbit anti-cleaved caspase 3 (CASP3, 1:500, 9661, Cell Signaling); rabbit anti-caveolin 1 (CAV1, 1:500, PA5-17447, Thermofisher); mouse anti-CD45 (CD45, 1:250, MA5-28384, Invitrogen); rabbit anti-claudin 10 (CLDN10, 1:500, 38-8400, Invitrogen); Alexa 488-conjugated mouse anti-claudin 5 (CLDN5, 1:250, 352588, Invitrogen); mouse anti-E-cadherin (ECAD, 1:500, 7D6, Developmental Studies Hybridoma Bank); mouse anti-E-cadherin clone 36 (ECAD, 1:500, 610181, BD Biosciences); rabbit anti-Avian erythroblastosis virus E-26 (v-ets) oncogene related (ERG, 1:250, ab92513, Abcam); rabbit anti-Keratin 14 (KRT14, 1:500, RB9020P0, Thermofisher); rabbit anti-NK2 homeobox 1 (NKX2-1, 1:500, AB133638, Abcam); mouse anti-NK2 homeobox 1 (NKX2-1, 1:500, TTF-1-L-CE, Leica Biosystems); Sambucus Nigra Lectin (SNA, EBL) Fluorescein (EB Lectin, 1:500, FL-1301-2, Vector Laboratories); Alexa488-conjugated mouse anti-ACTA2 (SMA, 1:1000, sc-32251 AF488, Santa Cruz Biotechnology); Cy3-conjugated mouse anti-ACTA2 (SMA, 1:1000, C6198, Sigma); Alexa647-conjugated mouse anti-ACTA2 (SMA, 1:1000, sc-32251 AF647, Santa Cruz Biotechnology); goat anti-SRY-box-containing gene 2 (SOX2, 1:250, AF2018, R&D Systems); rabbit anti-SRY-box-containing gene 2 (SOX2, 1:250, ab97959, Abcam); rabbit anti-SRY-box-containing gene 9 (SOX9, 1:1000, ab5536, Millipore); goat anti-SRY-box-containing gene 9 (SOX9, 1:500, AF3075, R&D Systems); goat anti-SRY-box-containing gene 17 (SOX17, 1:250, AF1924-SP, R&D Systems).

The following antibodies were used for the mouse data: rat anti-E-cadherin (ECAD, 1:1,000, 131900, Invitrogen); rabbit anti-Keratin 14 (KRT14, 1:500, RB9020P0, Thermofisher); goat anti-SRY-box-containing gene 2 (SOX2, 1:250, AF2018, R&D Systems); rabbit anti-SRY-box-containing gene 2 (SOX2, 1:250, ab97959, Abcam); goat anti-SRY-box-containing gene 9 (SOX9, 1:1,000, AF3075, R&D Systems), rabbit anti-SRY-box-containing gene 9 (SOX9, 1:1,000, AB5535, Millipore).

### Section and Whole-Mount Immunostaining

Immunostaining was performed similar to our mouse protocols previously published with minor modifications (Ostrin *et al*., 2018; Gerner-Mauro, Akiyama and Chen, 2020). Eggs of all ages were windowed, and embryos were euthanized by decapitation. Perfusion was performed by flushing the right ventricle of the heart with phosphate-buffered saline (PBS, pH 7.4). The lungs were dissected from the embryos and fixed with 0.5% paraformaldehyde (PFA; P6148, Sigma) in PBS for 3 hours on a rocker at room temperature. The lungs were washed at least one night in PBS at 4°C before processing. For section staining, either whole lungs or single lobes were cryoprotected with 20% sucrose with 1/10 volume optimal cutting temperature medium (OCT; 4583, Tissue-Tek) in PBS overnight at 4°C. The samples were embedded in OCT in embedding molds and sectioned at either 10, 20, or 30 µM onto Superfrost Plus slides (22-037-246, Thermofisher). Slides were washed with PBS three times at room temperature and were blocked in PBS 0.3% Triton X-100 (PBST) with 5% donkey serum (017-000-121, Jackson ImmunoResearch). Primary antibodies were diluted in PBST and incubated overnight at 4°C. Slides were washed in a coplin jar with PBS for 0.5 hours the next day. Secondary antibodies were diluted in PBST and added to slides and incubated at room temperature for 1.5 hours (1:1,000, Jackson ImmunoResearch or Invitrogen) and 4′,6-diamidino-2-phenylindole (DAPI, 0.5 μg/mL final concentration). Sections were washed again for 0.5 hours in PBS and mounted with Aquapolymount (18606, Polysciences).

For whole mount immunostaining, lungs were either processed as whole lungs, whole lobes, or as strips, cut from the morphological ridge from the first cranial costal sulci (Palmer and Nelson, 2020).Whole mount immunostaining was performed as previously published (Ostrin *et al*., 2018; Gerner-Mauro, Akiyama and Chen, 2020). Images were captured on an optical projection tomography (OPT) or an Olympus FV1000 confocal microscope.

### RNAscope Multiplex Fluorescent Staining

Lungs were excised from chicken embryos at the desired timepoint and snap frozen in OCT. 10 µM sections were collected on Superfrost Plus slides and the RNAscope® Multiplex Fluorescent Reagent Kit v2 assay was performed according to the manufacturer’s protocol for fresh frozen tissue (323100, Advanced Cell Diagnostics). Probes (Gg-*VEGFA*, 1095161-C1; Gg-*KRT14*, 1095211-C2; and Gg-*LAMP3*, 1095191-C3) were designed by the manufacturer and were screened for specificity using the RNAScope Hiplex platform (324409, Advanced Cell Diagnostics).

### OPT Microscopy

OPT microscopy was performed as previously published (Ostrin *et al*., 2018; Gerner-Mauro, Akiyama and Chen, 2020) with minor changes. Samples were washed and embedded in a 1% agarose gel (20-102GP, Genesee Scientific) dissolved in water. After curing, agarose gels were cut and mounted to the stages provided by the manufacturer (Bioptonics). Samples were dehydrated overnight with 100% methanol. The next day, samples were cleared in BABB (1:2 mixture of benzyl alcohol (24122, Sigma) and benzyl benzoate (B6630, Sigma), respectively) overnight. Samples were imaged once optically cleared. Reconstruction of the samples was performed using NRecon (BrukermicroCT, BE).

### Cell dissociation and fluorescence-activated cell sorting (FACS)

Lung dissociation was performed with some modifications to our previous publication (Cain, Hernandez and Chen, 2020). Whole lobes were harvested from E13 and E20 lungs, with four or two lobes pooled, respectively. Lobes were placed in Leibovitz’s Medium (21-083-027, Gibco), minced with forceps, and digested in Leibovitz’s with 2 mg/mL Collagenase Type I (LS004197, Worthington), 2 mg/mL Elastase (LS002294, Worthington), and 0.5 mg/mL DNase I (LS002007, Worthington) for 40 minutes at 37°C. Halfway into the digestion step, the tissue was pipetted for mechanical trituration. To stop the reaction, we added fetal bovine serum (FBS, 10082-139, Invitrogen) to a final concentration of 20%. We transferred the samples to ice and did the following steps at 4°C. The solution was mechanically homogenized with pipetting and then filtered with a 70 µm cell strainer (352350, Falcon). Samples were then centrifuged and washed with Leibovitz’s + 10% FBS, cells were resuspended in the same solution, and then filtered into a 5 mL tube with cell strainer cap (352235, Falcon). To enrich epithelial cells, cells were stained with an antibody against the extracellular domain of ECAD (Developmental Studies Hybridoma Bank, 7D6, 1:50) for 30 minutes, then centrifuged as above and washed with Leibovitz’s + 10% FBS. Alexa Fluor 647 AffiniPure donkey anti-mouse (715-605-151, Jackson ImmunoResearch) was added to a 1:250 concentration and incubated for 30 minutes and washed as previously described. All samples were resuspended in Leibovitz’s + 10% FBS, re-filtered, and stained with SYTOX Blue (S34857, Invitrogen,) before sorting. Sorting was performed on a BD FACSAria II Cell Sorter. Red blood cells were excluded based on size (>50 on the FSC-A gating), single cells were selected, and dead cells excluded. All ECAD-positive cells were collected from each sample, although other lung populations were still present. Single cell libraries were performed using either the Single Cell Multiome ATAC + Gene Expression kit (E13) or Chromium Single Cell Gene Expression Solution Platform (E20) (10x Genomics). 10X scRNA-seq and sc-multiome data are deposited at GSE264623.

### ScRNA-seq data analysis

scRNA-seq data analysis was performed as previously described with minor modifications. Raw data was processed using CellRanger (10X Genomics) and data was aligned using Galgal6. Only the RNA data of the E13 multiome was used for this study. Downstream normalization, aggregation of datasets, and differential gene expression analysis were performed with Seurat version 4.3.0.1 (Hao *et al*., 2021).

Chicken epithelial cells were computationally isolated using *CDH1* to isolate from mesenchymal (*COL3A1*), immune (*PTPRC*), and endothelial cells (*CDH5*). Lung epithelial cell types for progenitors, airway, AT1, and AT2 cells were assigned by conserved markers across mouse and chicken (Little *et al*., 2019; Gerner-Mauro, Akiyama and Chen, 2020). UMAPs were made with either Seurat or scCustom (Marsh, Salmon and Hoffman, 2023). Top genes from “FindAllMarkers” function in Seurat were plotted as heatmaps. Pseudotime trajectory analysis was performed with Monocle 3 (Cao *et al*., 2019). Modulescore was calculated using top 20 genes from AT1, AT2, and luminal cells. Volcano plot for luminal cells was plotted using EnhancedVolcano from values obtained from “FindMarkers” from aggregated luminal cells. GO Analysis on luminal cells was performed by taking the top 200 genes at either E13 or E20 and running these genes through DAVID (Sherman *et al*., 2022). Normalized counts of shared genes between mouse and chicken in pseudobulk SOX9, AT1, and AT2 cells were plotted as published (Travaglini *et al*., 2020). Ensembl IDs from uncharacterized genes were removed for GO analysis and psuedobulk plots. Mouse epithelial cells were used from our previously published datasets (Gerner-Mauro, Akiyama and Chen, 2020; Vila Ellis *et al*., 2020).

### Stereo-seq

E20 chicken lungs were harvested, and flash frozen in Tissue-Tek OCT medium in ethanol/dry ice bath, and then stored at −80°C until used for a Stereo-seq experiment (Chen *et al*., 2022, 2023; Jiang *et al*., 2024). Cryo-sections were cut at the thickness of 10 μm and mounted on Stereo-seq permeabilization chips (210CP118, STOmics) or transcriptomics chips (210CT114, STOmics). Tissue fixation and the following spatial transcriptomics procedure were performed according to the vendor’s manual and previous publications (2–3). Briefly, the tissue section on the Stereo-seq chip (1 cm x 1 cm) was incubated at 37°C for 5 min and subsequently fixed in pre-cold methanol (34860, Sigma, precooled 30 mins at −20°C) at −20°C. Once the fixation was completed, Stereo-seq chip was taken out to allow the residual methanol to dry out in a chemical hood. The tissue section on the chip was then stained with nucleic acid reagent (Invitrogen, Q10212, 0.5% v/v) for 5 min and subsequently washed with 0.1x SSC buffer (Ambion, AM9770; containing 0.05 U/ml RNase inhibitor). The Nuclei images were captured using a Zeiss Axio Scan Z1 microscope (at EGFP wavelength) and then followed by incubating tissue section in the permeabilization buffer (111KP118, STOmics) for 15 minutes at 37°C. Stereo-seq transcriptomics chip-captured RNAs from the permeated tissue were then reverse transcribed for 3 hours at 42°C. Following the tissue removal, the cDNAs were released from the chip using the transcriptomics reagent kit (111KT114, STOmics). After the cDNA yield was size-selected, amplified, and purified, the concentration was quantified by Qubit dsDNA HS assay kit (Q32854, Invitrogen). 20 ng of cDNA from each sample were used for library construction using the library preparation kit (111KL114, STOmics) and subsequently for DNB (DNA Nano Ball) generation. Finally, the DNBs were sequenced on the DNBSEQTM T7 sequencing platform (Complete Genomics, San Jose, USA) with 50 bp read1 and 100 bp read2 (1000028455, Complete Genomics).

The raw sequencing reads were mapped to chicken reference genome GRC7b using Stereo-seq Analysis Workflow (SAW_7.1) https://github.com/STOmics/SAW. The .cellbin.gef output file from SAW pipeline was further normalized, using Stereopy pipeline with default parameters https://github.com/STOmics/Stereopy. The neighborhood graph of cells was determined using the PCA representation of the expression matrix, and the graph was embedded into two dimensions using Uniform Manifold Approximation and Projection (UMAP). Next, the cells were clustered using leiden algorithm. The module scores of cell types identified in both spatial transcriptome and 10X scRNA-seq (re-aligned to the chicken reference genome GRC7b using the CellRanger software v6.0.0) were calculated using the function score_genes from tools module in Scanpy package (v1.9.6). The top 20 genes identified in the scRNA-seq analysis of E20 chicken lung were used for the module score calculation. Module scores were plotted using the function plot.umap and plot.embedding implemented in the Scanpy python package. Gene enrichment analysis of spatial transcriptome analysis was accomplished by plot.rank_genes_groups in the Scanpy package. The python version used for all analysis is 3.8.

## ACKNOWLEDGEMENTS

The University of Texas MD Anderson Cancer Center DNA Analysis Facility and Flow Cytometry and Cellular Imaging Core Facility are supported by the Cancer Center Support Grant (P30CA016672). This work was supported by the University of Texas MD Anderson Cancer Center Retention Fund, and National Institutes of Health R01HL130129 and R01HL153511 (JC) and R00HL155845 (LVE).

## AUTHOR CONTRIBUTIONS

KNGM, PYL, RAP, and JC designed research; KNGM, LVE, GW, RN, and JC performed research and analyzed data; KNGM and JC wrote the paper; all authors read and approved the paper.

## DATA AVAILABILITY

E13 and E20 chicken lung scRNA-seq data and E20 chicken lung Stereo-seq data are available at the NCBI Gene Expression Omnibus database (GEO) under GSE264623 and GSE274970, respectively. Custom scripts are available on GitHub-chenlab-cchmc/chicken-lung-airflow. Source quantification data is provided with this paper.

