## Supplemental Figures for "Morphogenic, molecular, and cellular adaptations for unidirectional airflow in the chicken lung"

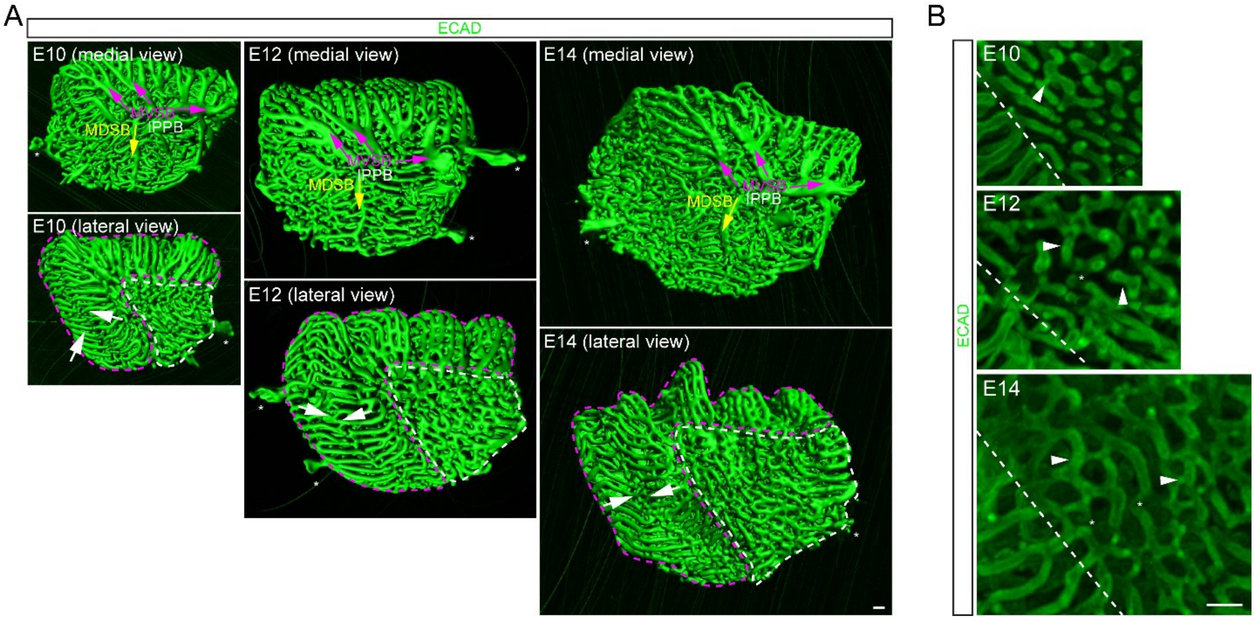


**Supplemental Figure 1: OPT imaging of paleopulmonic and neopulmonic regions of the chicken lung**

(A) OPT images of a whole lung to identify classic landmarks in paleopulmonic (magenta dash) and neopulmonic (white dash) regions. Neopulmonic regions likely correspond to primary termini that fuse within the anterior or posterior group. IPPB = intrapulmonary primary bronchus, MVSB = medioventral secondary bronchi, MDSB = mediodorsal secondary bronchi. Arrow: distal-long fusion between anterior and posterior groups. Asterisk: air sac. Scale bars: 250 μm.

(B) OPT images of intragroup distal-long (arrowhead) and proximal-short (asterisk) fusion. Dash: boundary between intergroup and intragroup parabronchi. Scale bars: 250 μm.

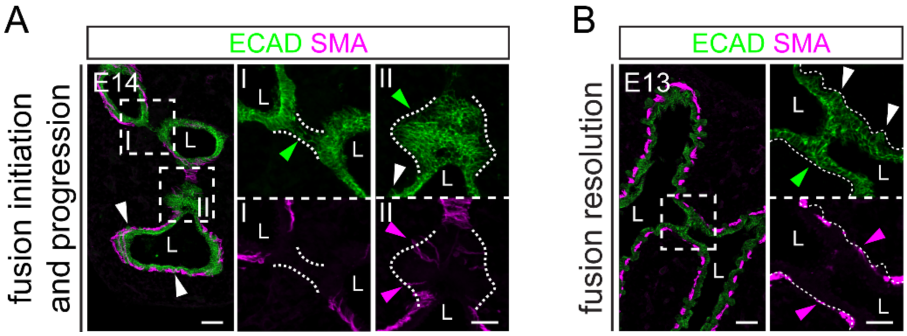


**Supplemental Figure 2: Neoform termini begin to form outside the fusion zone.**

(A) Confocal image of an E14 lung section showing initiation (I) and progression (II) phases of fusion (green arrowhead) with limited smooth muscle coverage (magenta arrowhead). White arrowhead: emerging neoform termini. L: lumen. Scale bars: 20 μm

(B) Confocal image of an E13 lung section showing neoform termini (white arrowhead) flanking a fused primary terminus that is resolving to connect the lumen (L) and is covered by smooth muscle (magenta arrowhead). Fusion is asynchronous and is more advanced for this region even at E13. Scale bars: 20 μm

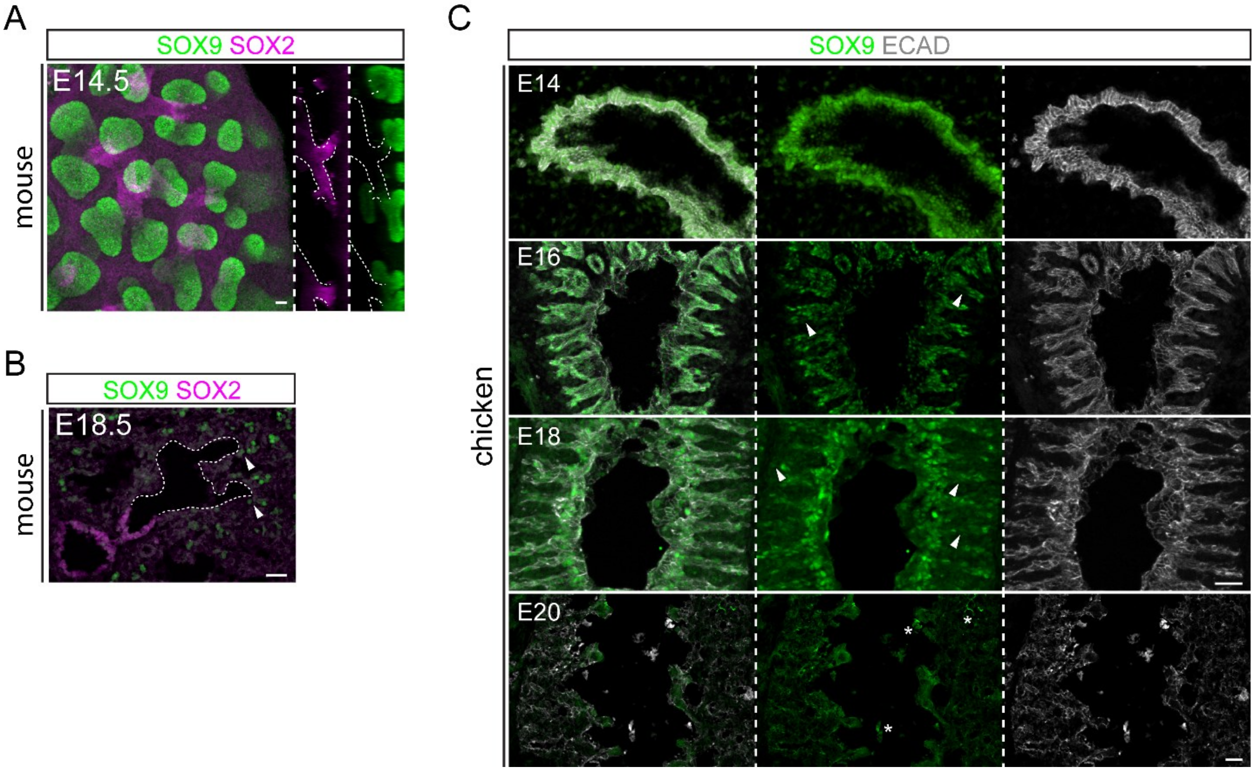


**Supplemental Figure 3: SOX9 downregulation during both mouse and chicken lung alveologenesis.**

(A) Confocal images of wholemount immunostained E14.5 mouse cranial lobe showing juxtaposition of distal (SOX9) and proximal (SOX2) markers. Scale bars: 20 µm.

(B) Confocal images of an E18.5 mouse lung section showing that SOX9 is still expressed in residual progenitors at branch tips (arrowhead) but downregulated in differentiating alveolar cells along alveolar ducts and alveolar sacs distal to SOX2+ airways (dash). Scale bars: 25 µm

(C) Confocal images of the chicken lung showing epithelial cells of radially growing neoform termini express SOX9 and only downregulate it during alveolar differentiation (arrowhead). Asterisk: background staining from tissue debris. Scale bars: 25 µm

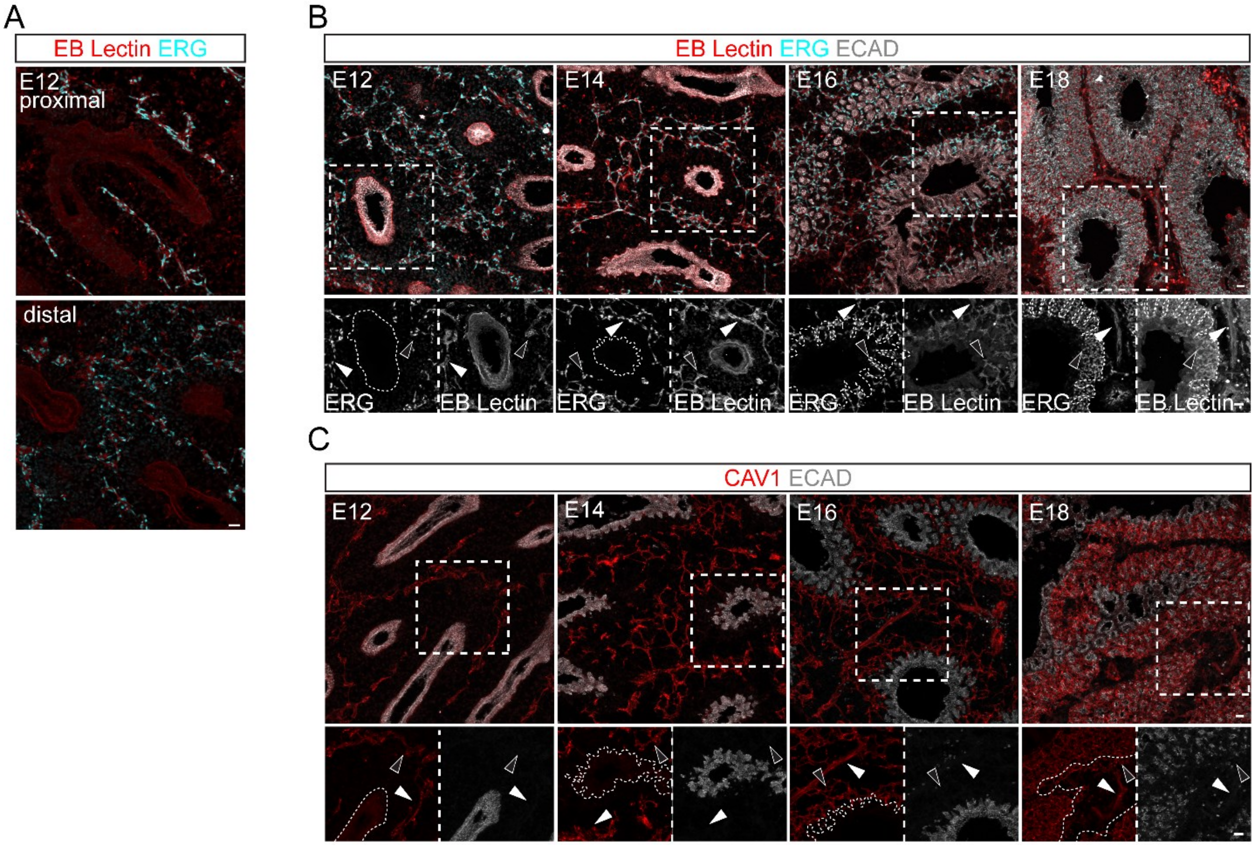


**Supplemental Figure 4: Additional evidence for primary plexus remodeling and secondary plexus formation.**

(A) Confocal images of E12 chicken lung sections showing tubular remodeled primary plexus (EB lection+ ERG+) in a proximal region before secondary plexus formation versus residual primary plexus capping branch tips in the distal region. Scale bars: 20 µm.

(B, C) Confocal images of chicken lung sections immunostained for vasculature (EB lectin; CAV1), vascular nuclei (ERG), and the epithelium (ECAD) over development, showing that the secondary plexus (open arrowhead) grows around the tubular macrovasculature (filled arrowhead) toward the epithelium (traced in insets). Scale bars: 10 µm.

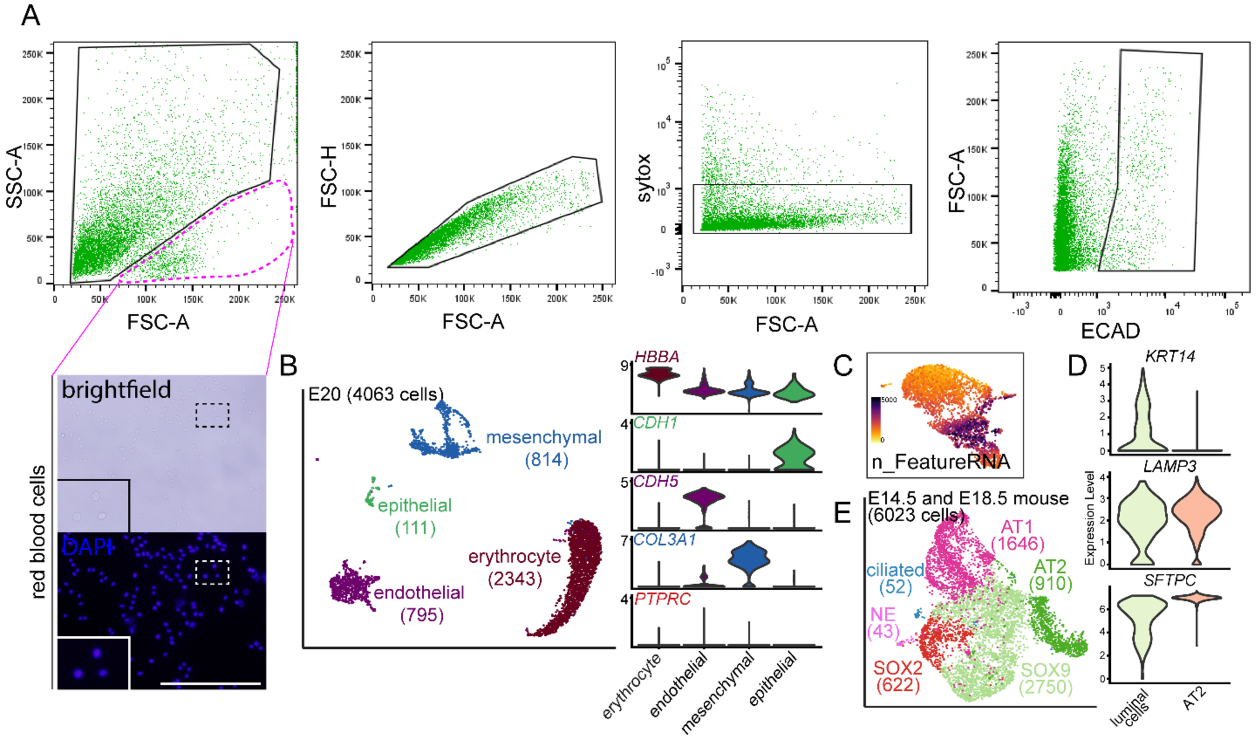


**Supplemental Figure 5: FACS gating of dissociated embryonic chicken lungs and additional analysis of single-cell profiling.**

(A) Representative FACS gating strategy to isolate ECAD+ chicken lung cells at E20. Red blood cells (erythrocytes) have large FSC-A (circle), as shown in the brightfield image of nucleated (DAPI) cells with their characteristic oval cell shape (inset). Scale bars: 50 µm.

(B) ScRNA-seq UMAP and corresponding violin plots of E20 CD45 depleted lungs highlighting the dominance of erythrocytes.

(C) Feature plot at E13 of transcript abundance (n_FeatureRNA).

(D) Violin plot comparison of AT2 and luminal cells showing specific expression of *KRT14*, but lower, variable expression of AT2 markers (*LAMP3* and *SFTPC*) in luminal cells.

(E) ScRNA-seq UMAPs of E14.5 and E18.5 mouse lung epithelial cells for comparison with E13 and E20 chicken data. NE: neuroendocrine cells.

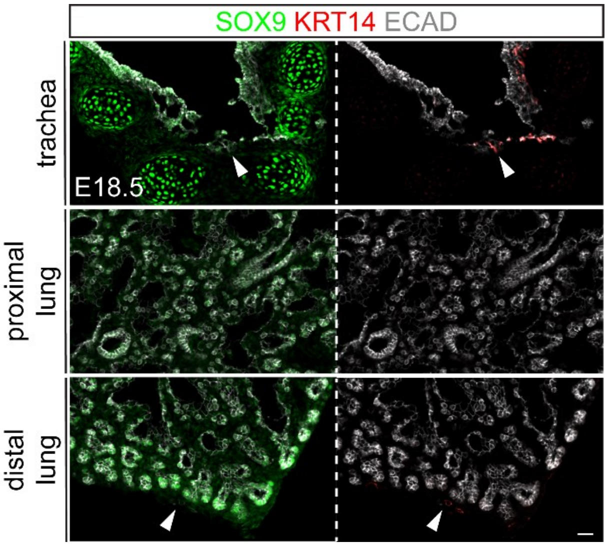


**Supplemental Figure 6: KRT14 is not expressed in alveolar epithelial cells in the mouse lung.**

Confocal images of the trachea, proximal, and distal lungs showing KRT14+ (arrowhead) basal cells surrounded by SOX9+ cartilage rings and mesothelial cells adjacent to residual SOX9+ epithelial progenitors. Scale bars: 25 μm.

**Supplemental Tables:**

**Table S1: Quantification of embryonic chicken lung branching and neoform termini formation (Fig. 2A, 2E).**

**Table S2: Differential gene expression analysis of E13 and E20 scRNA-seq for heatmaps (Fig. 5C, Fig. 6C), scatterplots (Fig. 5E, Fig 6E), and volcano plot (Fig. 6F).**

**Table S3: Cell type marker gene expression used for Stereo-seq (Fig. 8).**
